# Fluid amyloid-β (Aβ) biomarkers reflect early β-sheet-rich Aβ deposition during the preclinical stage in Alzheimer’s disease model 5XFAD mice

**DOI:** 10.64898/2026.04.06.716649

**Authors:** Hiroko Yagihara, Yuko Saito, Toshihide Takeuchi, Kazuhiko Seki, Eiko N. Minakawa

**Author notes:** **Correspondence Author: Eiko N. Minakawa, M.D., Ph.D.**, 4-1-1 Ogawa-Higashi, Kodaira, Tokyo, 187-8502, Japan.

## Abstract

Early detection of disease progression using clinically-relevant biomarkers in animal models is important for mechanistic studies and for developing therapeutics in neurodegenerative diseases including Alzheimer’s disease (AD). The preclinical stage of AD, when amyloid-β (Aβ) starts to accumulate before cognitive decline, provides a critical window for disease modification. In humans, decreases in cerebrospinal fluid (CSF) Aβ_42_ and the Aβ_42_/Aβ_40_ ratio in preclinical AD are considered to reflect the preferential sequestration of aggregation-prone Aβ_42_ into β-sheet-rich deposition in the brain, with corresponding changes being detectable in plasma. However, the extent to which these biomarker-pathology relationships are recapitulated in AD model mice remains incompletely defined. Here we show that CSF and plasma Aβ_42_ and the Aβ_42_/Aβ_40_ ratio decline with age in parallel with the progression of β-sheet-rich Aβ deposition in preclinical 5XFAD mice, one of the most widely used AD mouse models, as assessed through monthly profiling of these biomarkers. Notably, the CSF Aβ_42_/Aβ_40_ ratio showed a negative correlation with β-sheet-rich Aβ deposition in the brain, whereas CSF Aβ_40_ did not show a comparable association. In addition, the plasma Aβ_42_/Aβ_40_ ratio showed a positive correlation with the CSF Aβ_42_/Aβ_40_ ratio, suggesting that the plasma Aβ_42_/Aβ_40_ ratio may also reflect brain Aβ deposition in this model. The strength of these correlations differed by sex, suggesting that sex-dependent differences in the Aβ kinetics in this model may influence how closely fluid biomarkers reflect pathological progression. These findings support the potential utility of fluid Aβ as a pathology-linked, translatable biomarker in preclinical 5XFAD mice.

**Highlights:**

- Fluid Aβ biomarkers are associated with early Aβ deposition in preclinical 5XFAD mice.
- The CSF Aβ_42_/Aβ_40_ ratio negatively correlates with β-sheet-rich brain Aβ deposition.
- The plasma Aβ_42_/Aβ_40_ ratio positively correlates with the CSF Aβ_42_/Aβ_40_ ratio.
- Monthly profiling defines fluid Aβ biomarker dynamics in preclinical 5XFAD mice.
- Sex differences may affect biomarker-pathology relationships in these mice.

## Introduction

The preclinical stage of Alzheimer’s disease (AD) begins more than a decade before the onset of cognitive decline (Therriault et al., 2022). The initial AD pathology during the preclinical stage is the amyloid-β (Aβ) deposition in the senile plaques followed by hyperphosphorylated tau (p-tau) accumulation in the neurofibrillary tangles, eventually leading to neuronal dysfunction (Jack Jr. et al., 2024). Therapeutic interventions during the preclinical stage or early symptomatic stage offer an optimal opportunity to slow or halt disease progression before overt neuronal cell death, as recently confirmed by the efficacy of passive anti-Aβ immunotherapies in patients with early-phase AD (Heneka et al., 2024; Rafii and Aisen, 2023). Utilizing experimental animal models that recapitulate Aβ deposition in the preclinical stage could thus accelerate further development of therapeutic strategies against early AD to prevent or slow down the onset of cognitive decline (Drummond and Wisniewski, 2017).

A large number of mouse models of AD have been developed to recapitulate the preclinical stage of AD (Hashimoto and Saido, 2023). Age-dependent Aβ deposition before the onset of cognitive decline has been recapitulated in most of the transgenic mice expressing mutant *APP*, occasionally combined with mutant *PSEN1*, both of which are causative genes for familial AD (FAD) (Yokoyama et al., 2022). Early Aβ pathology before cognitive deficit has also been recapitulated in *App* knock-in mice with multiple causative gene mutations of FAD (Hashimoto and Saido, 2023; Saito et al., 2014).

To identify translatable therapeutic candidates using these preclinical AD model mice, precise detection of early Aβ deposition in mouse brains using methods applicable in clinical settings is essential (Jankowsky and Zheng, 2017). In patients with AD, positron emission tomography (PET) imaging with amyloid tracers provides a quantitative topographical assessment of brain Aβ plaque deposition. However, its widespread application in clinical settings is limited by specialized facility requirements and high cost for PET imaging and data analyses (Belder et al., 2023). Additionally, PET imaging of small mouse brains is limited by low spatial resolution, making it difficult to reliably detect Aβ plaque deposition (Frost et al., 2020).

Instead, recent technological advances in the detection of Aβ peptides in biofluids have established CSF and plasma biomarkers, including Aβ_42_ and the Aβ_42_/Aβ_40_ ratio, as practical alternatives that correlate with brain Aβ accumulation levels detected by amyloid PET (Jack Jr. et al., 2024). Aβ peptides are cleaved from amyloid beta precursor protein (APP) by the β- and γ-secretases into peptides of slightly varying length around 40 amino acids with different characteristics; Aβ_42_ is the most aggregation-prone species and the main constituent of brain Aβ plaque (DeTure and Dickson, 2019; Masters et al., 2015) The levels of CSF Aβ_42_ and the Aβ_42_/Aβ_40_ ratio were demonstrated to have high concordance with brain Aβ accumulation detected by amyloid PET or neuropathological analyses in humans (Hansson et al., 2018; Mattsson-Carlgren et al., 2022), and thus have been used in clinical practice in many countries (Hansson, 2021; Shaw et al., 2018). In addition, blood-based assays to detect these changes have been intensively developed considering the invasiveness of the CSF collection procedure (Hansson, 2021). The accuracy of plasma Aβ measurement has recently increased significantly, approaching that of CSF-based biomarkers with the development of automated ultrasensitive immunoassays (Jack Jr. et al., 2024). Therefore, validation of CSF and plasma Aβ biomarkers in mouse models of preclinical AD could facilitate therapeutic development by providing indicators of Aβ deposition in mouse brains during the preclinical stage that are translatable to human AD pathology.

Here, we investigated whether Aβ_42_ and the Aβ_42_/Aβ_40_ ratio in CSF could serve as indicators of preclinical brain Aβ deposition in 5XFAD mice, one of the most widely used AD models. 5XFAD mice express human *APP* carrying three mutations and human *PSEN1* carrying two mutations, all of which are causative for FAD, under the Thy1 promoter (Oakley et al., 2006). 5XFAD mice recapitulate the early and progressive Aβ deposition in the preclinical phase; Aβ plaques appear at two months of age in the brain and increase in an age-dependent manner, followed by neuronal loss beginning at four months and hippocampus-dependent cognitive deficits developing at six months of age (Oakley et al., 2006; Ohno et al., 2006). These findings suggest that the period corresponding to preclinical AD in 5XFAD mice is under six months of age. We therefore evaluated monthly changes in CSF biomarkers and assessed whether they correlated with brain Aβ deposition levels in these young preclinical 5XFAD mice. We also analyzed whether plasma Aβ levels correlate with CSF Aβ levels in preclinical 5XFAD mice to assess the utility of blood-based Aβ biomarkers during the preclinical stage.

## Results

### Aβ deposition progresses in an age-dependent manner in preclinical 5XFAD mouse brains

We first assessed Aβ deposition in the brains of 5XFAD mice at preclinical (1, 2, 3, 4, and 5 months of age) and advanced clinical (14 months of age) stages using immunohistochemistry with an anti-Aβ antibody. Consistent with previous studies (Oakley et al., 2006; Oblak et al., 2021), Aβ plaques were absent in the brains of 5XFAD mice at 1 month of age (data not shown), appeared at 2 months of age in both males and females, and showed an age-dependent increase during the preclinical stage (Supplementary Figures 1A and B). Consistent with previous findings, Aβ plaques developed earlier and faster in females than in males (Supplementary Figures 1A and B), a phenomenon attributed in previous studies to the estrogen response element (ERE) located upstream of the Thy-1 promoter in these mice (Javonillo et al., 2022; Sadleir et al., 2015).

We then performed thioflavin-S (thio-S) staining on the brain sections to investigate the deposition of Aβ with β-sheet-rich structure in 5XFAD mouse brains (Figures 1A and B). Thio-S selectively binds to β-sheet-rich structures and detects β-sheet-rich Aβ plaques that are also detectable with amyloid PET tracers (Ikonomovic et al., 2008; Jack Jr. et al., 2024). Thio-S-positive plaques were absent in the brains of 5XFAD mice at 1 month of age (data not shown), appeared at 2 and 3 months of age in female and male mice, respectively, and showed a significant age-dependent increase (Figures 1C and D).

**Figure 1.**
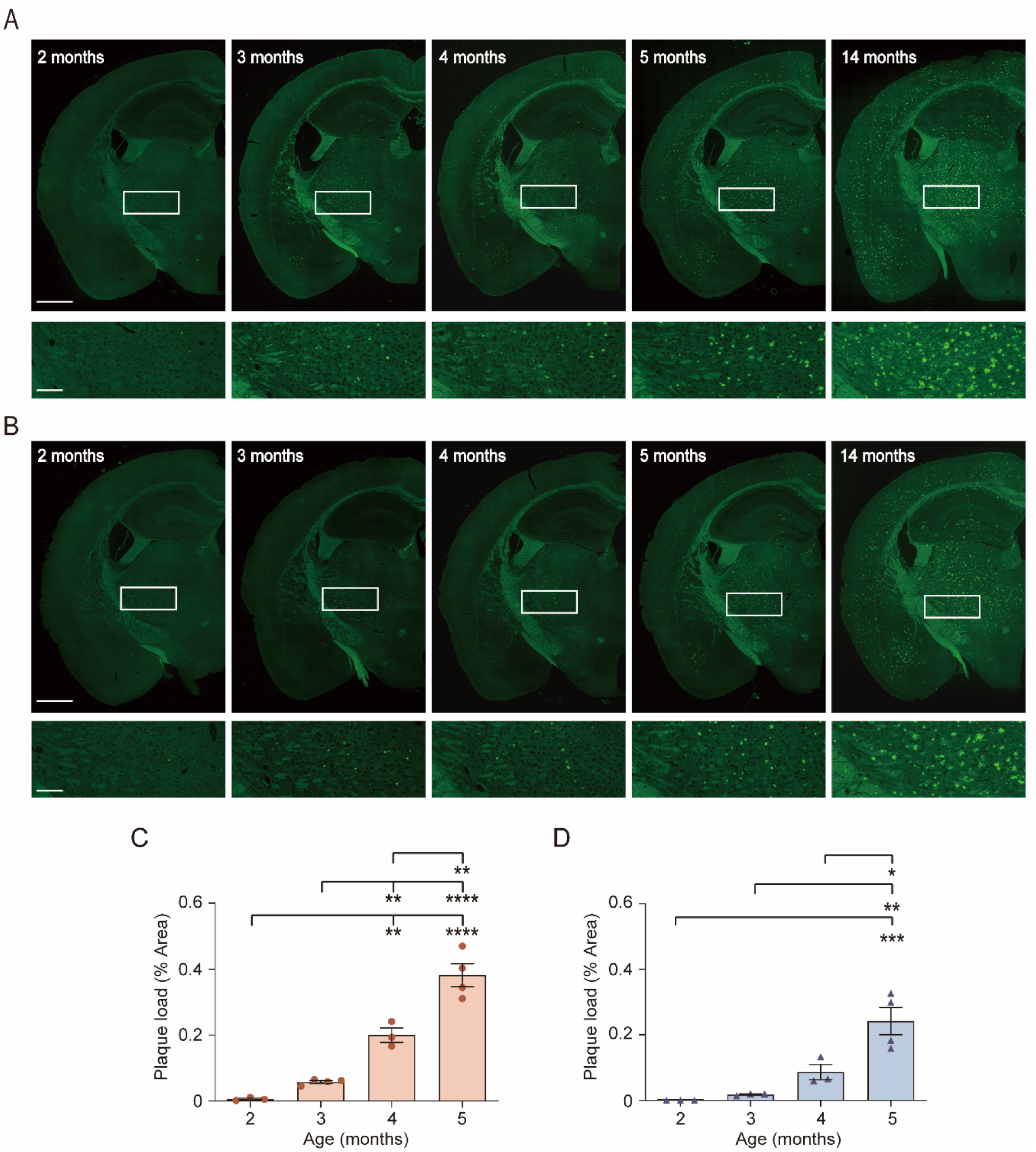
Monthly progression of β-sheet-rich Aβ deposition in preclinical 5XFAD mice. **(A and B)** Representative images of thioflavin-S staining in coronal brain sections of female (A) and male (B) 5XFAD mice in the preclinical (2–5 months old) and advanced clinical (14 months old) stages. The lower panels are magnified images of the insets in the upper panels. Scale bars, 1 mm (upper panels), 250 µm (lower panels). **(C and D)** Thioflavin-S-positive plaque load (total plaque area/total whole-brain surface area) in preclinical female (C) and male (D) 5XFAD mice. n = 3–4 mice for each group. Data are shown as mean ± S.E.M.; *p ≤ 0.05, **p ≤ 0.01, ***p ≤ 0.001, ****p ≤ 0.0001 in one-way ANOVA followed by Tukey’s multiple comparison test.

These findings confirm the age-dependent progression of Aβ deposition in the brains of preclinical 5XFAD mice.

### CSF Aβ_42_ and the Aβ_42_/Aβ_40_ ratio decrease in parallel with the progression of brain Aβ deposition in preclinical 5XFAD mice

A decrease in CSF Aβ_42_ levels and the resulting decline in the Aβ_42_/Aβ_40_ ratio have been established as one of the core biomarkers of AD, reflecting the accumulation of β-sheet-rich Aβ plaques in the brain in both preclinical and symptomatic phases of AD (Jack Jr. et al., 2024). Therefore, we next measured CSF Aβ_40_ and Aβ_42_ in 1- to 5-month-old 5XFAD mice to test whether the early decreases in CSF Aβ_42_ and the Aβ_42_/Aβ_40_ ratio during preclinical AD are recapitulated in preclinical 5XFAD mice. In both female and male mice, CSF Aβ_40_ remained unchanged with age (Figures 2A and 2B), which is consistent with observations in patients with preclinical AD (Hansson et al., 2019). In female mice, CSF Aβ_42_ and the CSF Aβ_42_/Aβ_40_ ratio decreased significantly with age (Figures 2C and E). In male mice, CSF Aβ_42_ and the CSF Aβ_42_/Aβ_40_ ratio displayed a tendency to decline in an age-dependent manner without reaching statistical significance (Figures 2D and F), which may be attributed to milder progression of Aβ accumulation in the brains of male 5XFAD mice than female mice (Figures 1C and D).

**Figure 2.**
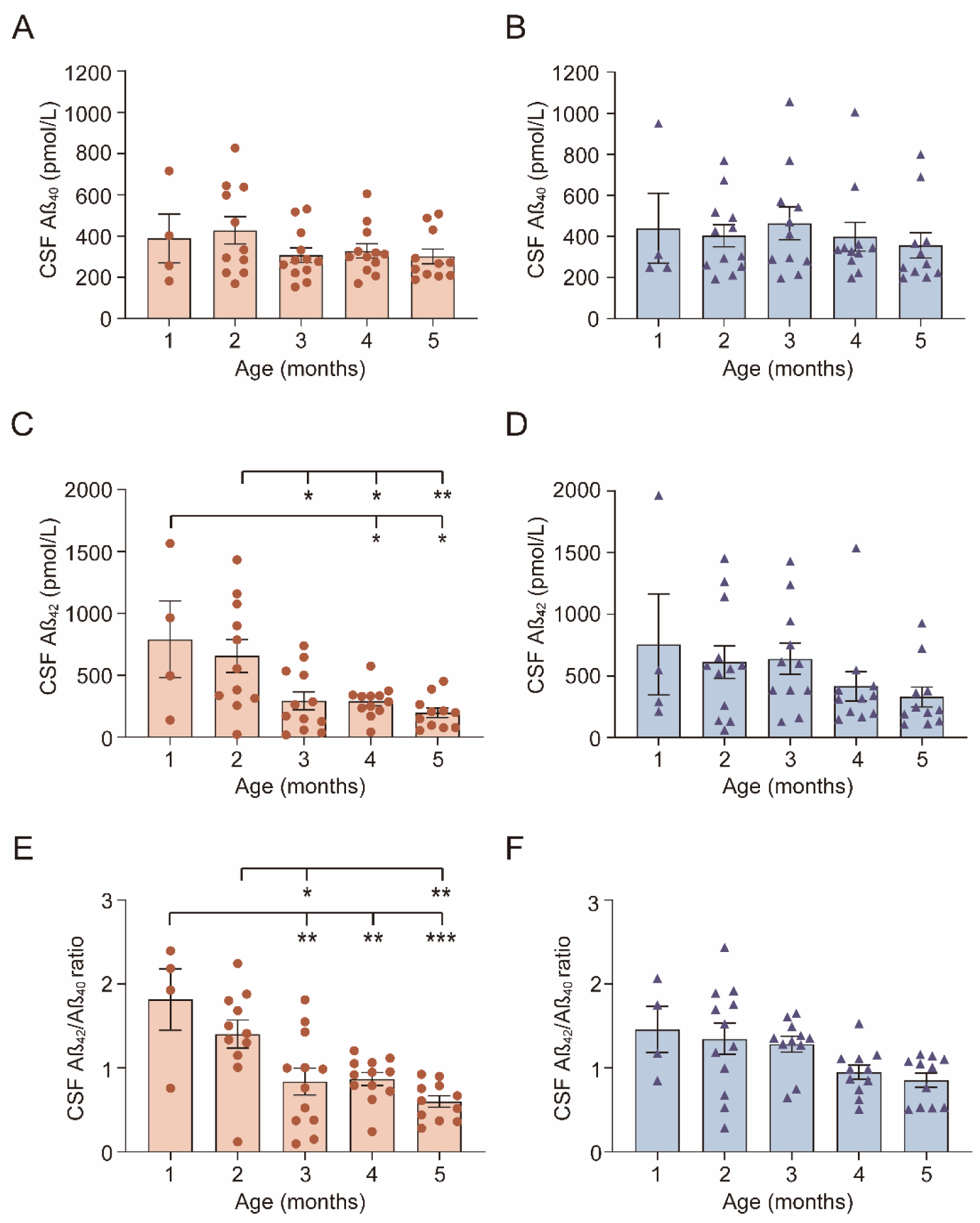
Age-dependent changes in CSF Aβ biomarkers align with progression of β-sheet-rich Aβ deposition in preclinical 5XFAD mice. CSF Aβ_40_ (A and B) and Aβ_42_ (C and D) levels and the Aβ_42_/Aβ_40_ ratio (E and F) in female (A, C, E) and male (B, D, F) preclinical 5XFAD mice (1–5 months old). n = 4–12 mice for each group. Data are shown as mean ± S.E.M; *p ≤ 0.05, **p ≤ 0.01, ***p ≤ 0.001 in one-way ANOVA followed by Tukey’s multiple comparison test.

We then analyzed the relationship between CSF Aβ and brain Aβ deposition in 5XFAD mice. In both male and female mice, CSF Aβ_40_ did not show a strong correlation with the thio-S-positive Aβ deposition in the brain (Figures 3A and B; r = − 0.174, p = 0.553 in females; r = − 0.432, p = 0.246 in males), as expected from the aforementioned age-dependent analyses (Figures 2A and B). In contrast, both CSF Aβ_42_ and the Aβ_42_/Aβ_40_ ratio showed stronger negative correlations than CSF Aβ_40_ with the thio-S-positive Aβ deposition in mouse brains (Figures 3C–F; for Aβ_42,_ r = − 0.453, p = 0.104 in females, r = − 0.587, p = 0.0963 in males; for Aβ_42_/Aβ_40,_ r = − 0.484, p = 0.0795 in females, r = − 0.878, p = 0.0018 in males). These results indicate that both CSF Aβ_42_ and the Aβ_42_/Aβ_40_ ratio reflect the β-sheet-rich Aβ deposition in the brains of preclinical 5XFAD mice better than CSF Aβ_40_, as in humans. Intriguingly, stronger correlations between the CSF Aβ_42_/Aβ_40_ ratio or Aβ_42_ and brain Aβ deposition were observed in males than in females (Figures 3C–F). In addition, the correlation between the CSF Aβ_42_/Aβ_40_ ratio and brain Aβ deposition was stronger than that between CSF Aβ_42_ and brain Aβ deposition, which reached statistical significance in males (Figures 3C–F). These results suggest that, even when disease progression is relatively mild, such as in preclinical male 5XFAD mice, the CSF Aβ_42_/Aβ_40_ ratio may reflect changes in brain Aβ deposition more precisely than CSF Aβ_42_ alone in 5XFAD mice.

**Figure 3.**
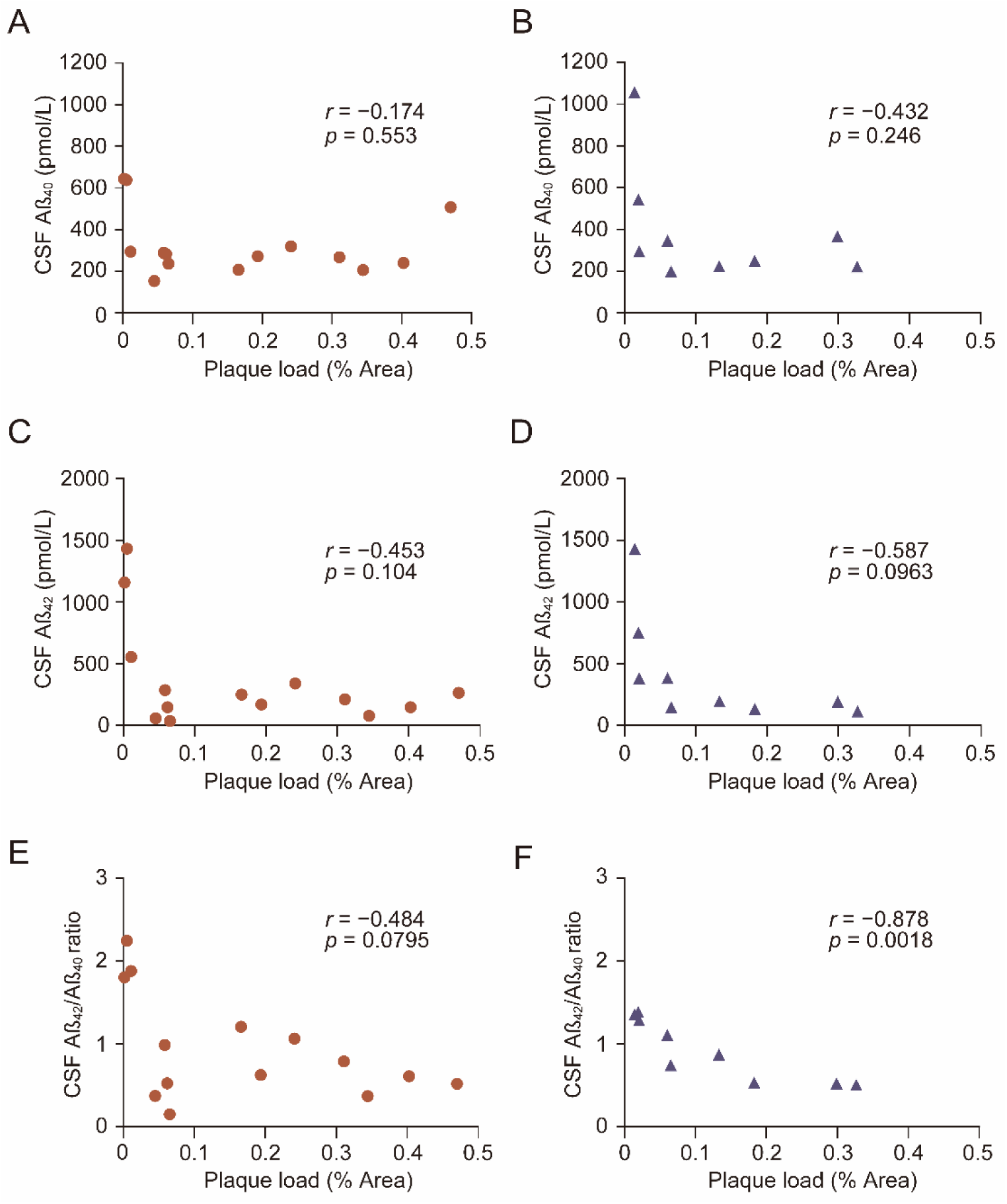
CSF Aβ_42_ and the CSF Aβ_42_/Aβ_40_ ratio correlates with β-sheet-rich Aβ deposition in preclinical 5XFAD mice. Scatter plots of thioflavin-S-positive plaque load and CSF Aβ_40_ (A and B) or Aβ_42_ (C and D) levels or the Aβ_42_/Aβ_40_ ratio (E and F) in female (A, C, E) and male (B, D, F) preclinical 5XFAD mice (1–5 months old). Females aged 2–5 months, n=14; males aged 3–5 months, n=10. Pearson’s correlation analysis was used to calculate *r* and p values for all scatter plots.

### Plasma Aβ_42_ and the Aβ_42_/Aβ_40_ ratio decrease with age in preclinical 5XFAD mice

Plasma biomarkers are increasingly being used to detect early AD-related changes because plasma can be collected more easily and less invasively than CSF. Previous studies have demonstrated decreased plasma Aβ_42_ levels and the Aβ_42_/Aβ_40_ ratio in patients with AD including those in the preclinical phase (Cheng et al., 2022; Salvadó et al., 2023; Trelle et al., 2025). We therefore quantified plasma Aβ_40_ and Aβ_42_ in 5XFAD mice aged 1 – 5 months to determine whether plasma Aβ biomarkers reflect the age-dependent brain Aβ deposition in preclinical 5XFAD mice. In both females and males, plasma Aβ_40_ showed no age-dependent change (Figures 4A and B), as expected from the aforementioned CSF measurement (Figures 2A and B). In contrast, both plasma Aβ_42_ and the Aβ_42_/Aβ_40_ ratio decreased with age (Figure 4C–F). Notably, in females, the decrease in plasma Aβ_42_ from the baseline (1 month of age) reached statistical significance at an earlier timepoint than CSF Aβ_42_ (Figures 2C and 4C; 3 months of age in plasma; 4 months of age in CSF). In addition, in males, the decrease in plasma Aβ_42_ and the Aβ_42_/Aβ_40_ ratio from the baseline (1 – 2 months of age) reached statistical significance at 4 or 5 months of age (Figures 4D and F). This was in contrast with the changes in CSF Aβ_42_ and the Aβ_42_/Aβ_40_ ratio, both of which did not reach statistical significance in males (Figures 2D and F). These observations suggest that the plasma Aβ_42_ and the Aβ_42_/Aβ_40_ ratio may detect brain Aβ deposition with higher sensitivity than CSF Aβ_42_ or the Aβ_42_/Aβ_40_ ratio in preclinical 5XFAD mice.

**Figure 4.**
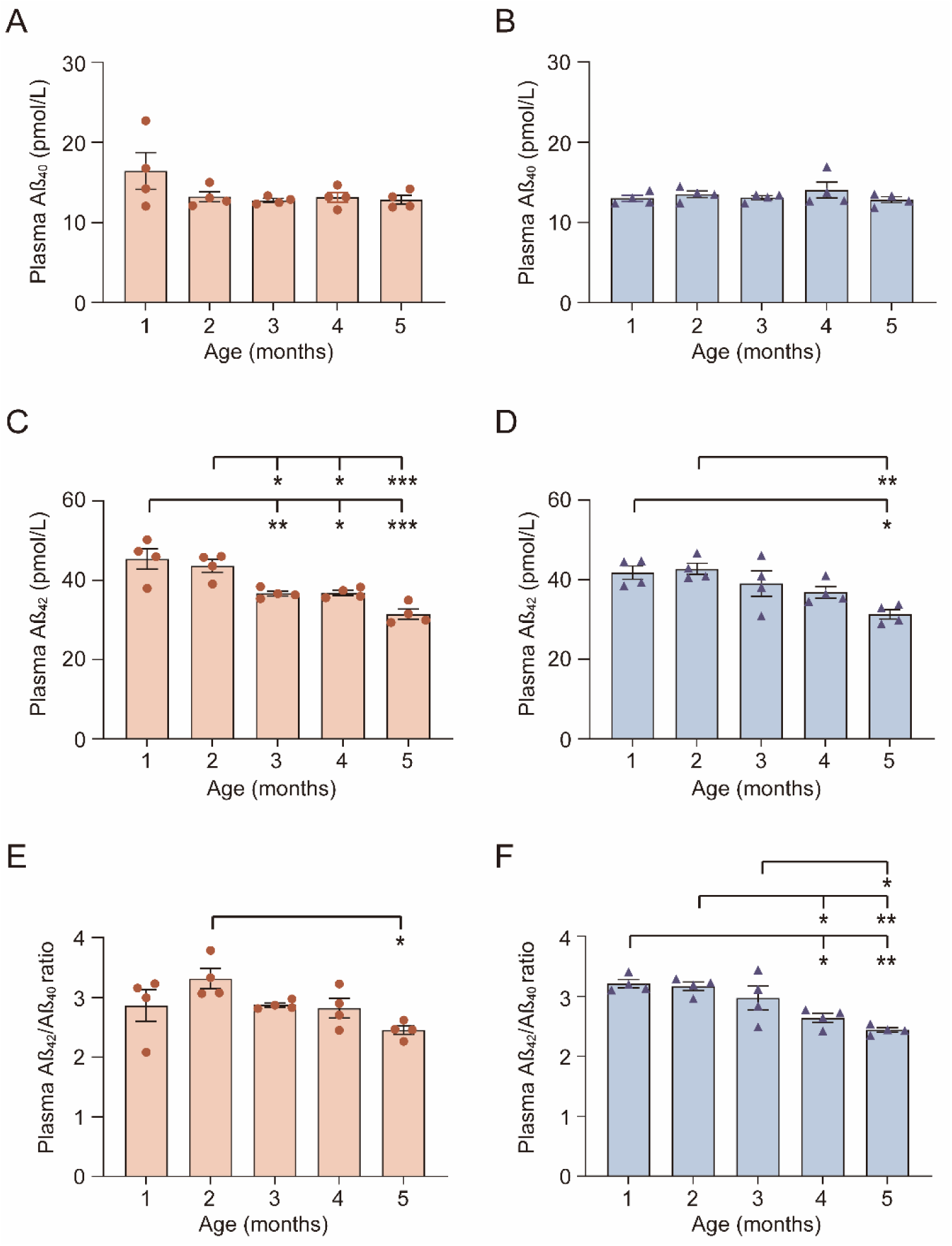
Age-dependent changes in plasma Aβ biomarkers align with progression of β-sheet-rich Aβ deposition in preclinical 5XFAD mice. Plasma Aβ_40_ (A and B) and Aβ_42_ (C and D) levels and the Aβ_42_/Aβ_40_ ratio (E and F) in female (A, C, E) and male (B, D, F) preclinical 5XFAD mice (1–5 months old). n = 4 mice for each group. Data are shown as mean ± S.E.M.; *p ≤ 0.05, **p ≤ 0.01, ***p ≤ 0.001 in one-way ANOVA followed by Tukey’s multiple comparison test.

### Relationships between plasma and CSF support the plasma Aβ_42_/Aβ_40_ ratio as a potential indicator of brain Aβ deposition in preclinical 5XFAD mice

We further examined whether plasma Aβ_42_ and the Aβ_42_/Aβ_40_ ratio correlated with their CSF counterparts in preclinical 5XFAD mice. In both females and males, Aβ_40_ showed no correlation between CSF and plasma (Figures 5A and B; r = − 0.150, p = 0.529 in females; r = − 0.218, p = 0.357 in males). In females, plasma and CSF showed positive correlations in both Aβ_42_ and the Aβ_42_/Aβ_40_ ratio, with a stronger correlation for the Aβ_42_/Aβ_40_ ratio that reached statistical significance (Figures 5C and E; for Aβ_42,_ r = 0.348, p = 0.132; for Aβ_42_/Aβ_40,_ r = 0.548, p = 0.0123). In males, plasma and CSF Aβ_42_ did not correlate, whereas the plasma and CSF Aβ_42_/Aβ_40_ ratio showed a modest positive correlation (Figures 5D and F; for Aβ_42,_ r = 0.0654, p = 0.784; for Aβ_42_/Aβ_40,_ r = 0.399, p = 0.0817). Considering that the CSF Aβ_42_/Aβ_40_ ratio negatively correlated with thio-S–positive brain Aβ with statistical significance in both female and male preclinical 5XFAD mice (Figure 3C–F), these findings suggest that the plasma Aβ_42_/Aβ_40_ ratio may provide an estimate of early brain Aβ deposition that aligns with CSF changes in preclinical 5XFAD mice.

**Figure 5:**
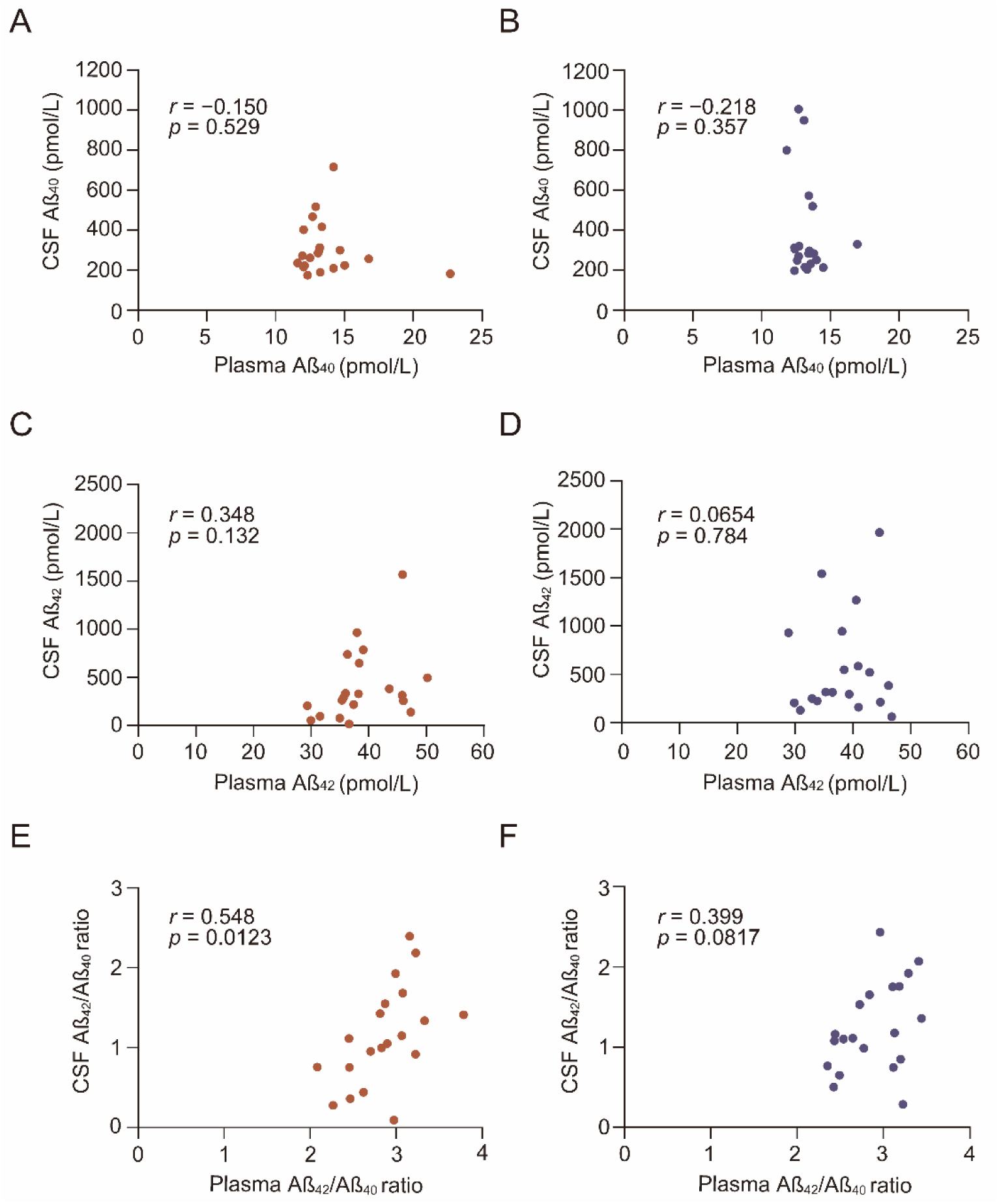
The plasma Aβ_42_/Aβ_40_ ratio correlates with the CSF Aβ_42_/Aβ_40_ ratio in preclinical 5XFAD mice. Scatter plot of plasma Aβ_40_ (A and B), Aβ_42_ (C and D), and the Aβ_42_/Aβ_40_ ratio (E and F) with their CSF counterparts in female (A, C, and E) or male (B, D, and F) preclinical 5XFAD mice (1–5 months old). n=20 for females and males, respectively. Pearson’s correlation was used to calculate *r* and p values for all scatter plots.

## Discussion

AD is characterized by a long preclinical phase during which Aβ pathology accumulates before the onset of overt cognitive decline. In the present study, we investigated whether fluid Aβ biomarkers reflect early β-sheet-rich Aβ deposition in the brain during the preclinical stage in 5XFAD mice. We found that CSF Aβ_42_ and the CSF Aβ_42_/Aβ_40_ ratio showed negative correlations with thio-S–positive Aβ deposition in the brain, whereas CSF Aβ_40_ remained unchanged. Plasma Aβ_42_ and the plasma Aβ_42_/Aβ_40_ ratio decreased in an age-dependent manner during the preclinical stage, and the plasma Aβ_42_/Aβ_40_ ratio correlated positively with its CSF counterpart. Together, these findings indicate that young 5XFAD mice reproduce key changes in Aβ-related fluid biomarkers observed in human preclinical AD, and suggest that this model may serve as an animal model of preclinical AD for evaluating candidate therapeutics targeting Aβ pathology by using fluid Aβ biomarkers as translational indicators of therapeutic efficacy.

A major strength of the present study is that it characterizes biofluid Aβ dynamics across the early preclinical window in 5XFAD mice with monthly resolution and relates these changes directly to histologically detected β-sheet-rich Aβ deposition. Although 5XFAD mice are widely used in AD research, relatively few studies have focused specifically on fluid biomarker changes during the preclinical phase, when disease-modifying interventions are expected to be most effective. Andersson et al. reported a decline in the CSF Aβ_42_/Aβ_40_ ratio during the preclinical stage of 5XFAD mice and identified soluble brain protofibril composition as a key predictor of CSF biomarker changes (Andersson et al., 2025), while Kwak et al. demonstrated decreases in CSF and plasma Aβ_42_ at a more advanced stage (Kwak et al., 2020). Similar preclinical reductions in CSF and/or blood Aβ_42_-related measures have been reported in several other widely used mouse models of AD, including *App* knock-in and *APP*/*PSEN1*-related lines (Andersson et al., 2025, 2023; Izco et al., 2014; Kawarabayashi et al., 2001; Maia et al., 2013). However, studies that assess the early preclinical phase with monthly resolution, analyze both CSF and plasma/serum Aβ measures in parallel, and directly compare these fluid biomarker changes with brain Aβ deposition appear to be limited in AD model mice, including 5XFAD mice. While previous studies support the general concept that early fluid biomarker changes are reproducible across multiple AD model mice, these studies also indicate that the timing and magnitude of fluid Aβ biomarker changes and their relationships with brain Aβ deposition in these AD model mice depend largely on the phenotypic characteristics of each mouse model, including the progression rate of brain Aβ deposition, and also on the assay platform used for Aβ detection. In the present study, we detected significant declines in CSF and plasma Aβ_42_ levels and the CSF Aβ_42_/Aβ_40_ ratio from baseline at 3 months of age in female 5XFAD mice, which, to the best of our knowledge, represents the earliest timepoint reported to date. These changes were detectable with conventional ELISA, which is more cost-effective than the recently developed ultrasensitive immunoassays. Our study therefore extends prior work and highlights the utility of 5XFAD model mice as a preclinical AD model by providing a more temporally refined characterization of the preclinical stage, assessing both CSF and plasma Aβ in parallel, and relating these changes to thio-S–positive β-sheet-rich Aβ deposition in the brain. Since thio-S-positive plaques are closely related to the species detected by amyloid PET in humans (Ikonomovic et al., 2008), our findings also enhance the translational relevance of fluid Aβ biomarkers in 5XFAD mice.

The observed decline in CSF Aβ_42_ and the CSF Aβ_42_/Aβ_40_ ratio in correlation with thio-S-positive Aβ deposition (Figures 3C–F) and the absence of change in CSF Aβ_40_ alone are consistent with the general interpretation of these biomarkers in human AD. Because Aβ_42_ is more aggregation-prone than Aβ_40_ and is preferentially incorporated into insoluble amyloid deposits, the reduction in these CSF measures likely reflects sequestration of soluble Aβ_42_ into progressively accumulating brain aggregates (Gravina et al., 1995; Iwatsubo et al., 1994; Jarrett et al., 1993; Strozyk et al., 2003). The stronger association of the CSF Aβ_42_/Aβ_40_ ratio with brain Aβ deposition, compared with CSF Aβ_42_ alone, also mirrors clinical findings showing the utility of the Aβ_42_/Aβ_40_ ratio rather than Aβ_42_ alone (Frisoni et al., 2025).

An additional important finding is that plasma Aβ_42_ and the plasma Aβ_42_/Aβ_40_ ratio decreased in an age-dependent manner during the preclinical stage in 5XFAD mice and appeared to show clearer changes than their CSF counterparts, particularly in males (Figure 4C–F). In humans, CSF Aβ_42_ or Aβ_42_/Aβ_40_ ratio has traditionally been considered more tightly linked to brain amyloid pathology than plasma Aβ_42_ or Aβ_42_/Aβ_40_ ratio, especially when assessed by conventional ELISA (Hansson, 2021). Several factors may account for this apparent difference in the mouse model. First, 5XFAD mice are an overexpression model with high production of human Aβ (Oakley et al., 2006), potentially generating a stronger peripheral signal in the plasma than is observed in sporadic human AD. Second, the rapid progression of Aβ plaque formation in 5XFAD mice may produce steep biomarker changes, allowing plasma changes to become detectable within a narrow time window, especially due to the smaller total plasma volume in mice. Third, since CSF samples are more technically challenging to collect than blood and have much smaller sample volumes, CSF Aβ measurement may result in greater variability due to pre-analytical factors, including collection efficiency, Aβ adsorption to collection tube walls, or blood contamination (Bjerke et al., 2010). These considerations suggest that the apparent sensitivity of plasma in this study does not indicate that plasma is superior to CSF for reflecting AD pathology in general. Rather, our data suggest that, in this specific model and assay condition, plasma Aβ_42_ and the plasma Aβ_42_/Aβ_40_ ratio may serve as practical surrogate indicators of early Aβ deposition.

The sex-related differences observed in this study also merit consideration. As reported previously, female 5XFAD mice developed Aβ pathology earlier and more rapidly than males (Figures 1C and D), likely due at least in part to the estrogen response element within the Thy1 promoter driving transgene expression (Oakley et al., 2006; Sadleir et al., 2015). Interestingly, however, correlations between CSF Aβ_42_-related measures and brain Aβ deposition were stronger in males despite their milder pathology (Figure 3C–F). The slower progression of amyloid deposition in males may create a broader detection window, enabling biofluid biomarkers to reflect a milder increase in brain Aβ deposition in a more gradual and linear manner. In contrast, the more rapid accumulation of brain Aβ in females may narrow this detection window, making cross-sectional correlations weaker. These findings suggest that sex influences not only the extent of amyloid deposition but also the temporal detectability of fluid biomarker changes in 5XFAD mice.

This study has several limitations. First, 5XFAD mice are a transgenic model harboring multiple mutations in *APP* and *PSEN1* and do not fully recapitulate the pathomechanisms of sporadic AD. Second, brain Aβ deposition was quantified using thio-S alone, without parallel Aβ immunohistochemistry or biochemical measurement of soluble and insoluble Aβ in the brain homogenates. Third, all analyses were performed cross-sectionally rather than longitudinally within the same animals. Fourth, as mentioned above, pre-analytical variability may influence absolute Aβ measurements, particularly in the CSF. Finally, this study focused on early Aβ-related changes and did not assess downstream events in the disease course of AD such as p-tau accumulation, neuroinflammation, or neurodegeneration.

Despite these limitations, the present study provides a temporally refined characterization of fluid Aβ biomarker dynamics during the preclinical stage of 5XFAD mice and demonstrates that these changes are linked to early β-sheet-rich brain Aβ deposition. Among the Aβ-related biomarkers examined, the Aβ_42_/Aβ_40_ ratio appeared to be the most consistent indicator of brain Aβ deposition, and plasma biomarkers may offer a practical complement to CSF biomarkers in this model. These findings, together with prior evidence that 5XFAD mice lack insertional mutagenesis and that their brain transcriptome overlaps with that of human AD (Goodwin et al., 2019; Preuss et al., 2020; Zhong et al., 2024), support the use of young 5XFAD mice as a model of preclinical AD for evaluating the efficacy of therapeutic interventions targeting early Aβ pathology. Future studies are warranted to determine whether longitudinal changes in fluid Aβ biomarkers can be tracked within the same animals, especially using plasma-based ultrasensitive immunoassays, and also whether these biomarkers respond to therapeutic interventions that slow or halt Aβ deposition. By enabling therapeutic effects to be assessed before overt cognitive decline, this mouse model may help accelerate the development of disease-modifying therapies that delay or prevent dementia.

## Materials and Methods

### Animals

All animal experiments were approved by the Animal Ethics Committee of the National Institute of Neuroscience, National Center of Neurology and Psychiatry (NCNP), Japan. 5XFAD mice (B6SJL-Tg(APPSwFlLon,PSEN1*M146L*L286V)6799Vas/Mmjax) were obtained from the Jackson Laboratory (Bar Harbor, ME, USA) and maintained by mating on a C57BL/6J background under a 12-hour light/dark cycle with food and water provided *ad libitum*.

### CSF, plasma, and brain collection

1, 2, 3, 4, 5, and 14-month-old 5xFAD mice were anesthetized by intraperitoneal administration of ketamine/xylazine mixture. CSF was collected between 9 a.m. and 12 p.m. from the cisterna magna using a glass capillary tube (1B150F-4; World Precision Instruments, Sarasota, FL, USA) as described previously (Liu and Duff, 2008). The collected CSF was centrifuged at 6,200 × g for 5 minutes at 4°C, and the supernatant was stored at −80°C until use. The pellet was used to assess blood contamination as described previously (Barten et al., 2011), and blood-contaminated CSF was excluded from further analyses. Whole blood was collected through cardiac puncture from the left ventricle into tubes containing ethylenediaminetetraacetic acid dipotassium salt (EDTA-2K) and centrifuged at 2,000 × g for 5 minutes at room temperature. The supernatant was collected and stored at −80°C until the time of assay.

For pathological evaluation, mice were deeply anesthetized and transcardially perfused with phosphate-buffered saline (PBS) followed by 10% neutral-buffered formalin (NBF). Brains were dissected out, post-fixed with 10% NBF for 48 hours at 4°C, transferred to PBS, and incubated at 4°C for a minimum of 24 hours before being embedded in paraffin and coronally sectioned at a thickness of 5 μm for further analysis.

### Thioflavin-S staining and immunohistochemistry

Deparaffinized sections were incubated with 1% (wt/vol.) thioflavin S (T1892, Sigma Aldrich St. Louis, MO, USA) in distilled water for 8 minutes at room temperature in the dark and rinsed in 80% ethanol, 95% ethanol, and distilled water as described previously (Yan et al., 2020). The sections were examined with a fluorescence microscope (BZ-X710, Keyence, Osaka, Japan), and the thio-S-positive plaques in the whole brain section were quantitatively analyzed as described previously using dedicated software (Hybrid Cell Count, Keyence, Osaka, Japan) (Danhash et al., 2025; Minakawa et al., 2017). Immunohistochemistry for Aβ plaques was performed as described previously (Minakawa et al., 2017) with slight modification. Deparaffinized sections were autoclaved in citrate buffer (pH 6.8) at 105°C for 10 min followed by 100% formic acid treatment at room temperature for 10 min for antigen retrieval. After endogenous peroxidase deactivation and blocking, sections were incubated with mouse monoclonal anti-β-amyloid antibody (clone 4G8; 800703; BioLegend, San Diego, CA, USA) at 4°C overnight. Then the sections were incubated with Histofine Simple Stain MAX PO (MULTI) (424152; Nichirei Biosciences Inc., Tokyo, Japan) for 30 min at room temperature, colorized with 3,3’-diaminobenzidinetetrahydrochloride (DAB) solution containing 0.03% hydrogen peroxide, and counterstained with hematoxylin. The sections were examined with bright-field microscopy (BZ-X710, Keyence, Osaka, Japan).

### Aβ enzyme-linked immunosorbent assay (ELISA)

CSF and plasma were thawed on ice and centrifuged at 6,200 × g for 5 minutes at 4 °C. The supernatant was collected and used immediately for ELISA. Aβ_40_ and Aβ_42_ concentrations were measured using a human β-Amyloid ELISA kit (298–64601 and 296–64401; Fujifilm Wako Pure Chemical Corporation, Osaka, Japan) according to the manufacturer’s instructions. Sample dilution fold for each analyte was as follows: for Aβ_40_, 33 to 50-fold for CSF and 4-fold for plasma; for Aβ_42_, 100-fold for CSF and 20-fold for plasma.

### Statistical analysis

Monthly changes in the thio-S-positive plaques and CSF and plasma Aβ were analyzed by one-way analysis of variance (ANOVA) followed by Tukey’s multiple comparisons test. Correlations between thio-S accumulation in the brain and CSF Aβ biomarkers, as well as CSF and plasma Aβ biomarkers, were examined using Pearson’s correlation analysis. All statistical analyses were performed using GraphPad Prism 9 (Dotmatics, Boston, MA, USA). P < 0.05 was considered to indicate a statistically significant difference.

## Author Contributions

Conceptualization: E.N.M., T.T.; Investigation and Formal analysis: H.Y., E.N.M.; Funding acquisition: Y.S., T.T., K.S., E.N.M.; Visualization: H.Y., E.N.M.; Supervision: Y.S., K.S., E.N.M.; Writing – original draft: H.Y., E.N.M.; Writing – review and editing: H.Y., Y.S., T.T., K.S., E.N.M.

## Acknowledgements

The authors are grateful to Ms. Hiromi Fujita, Yoshiko Hara, and Hisae Kikuchi for their technical support.

## Data Availability

The data that support the findings of this study are available from the corresponding author, upon reasonable request.

---

**Supplementary Figure 1.**
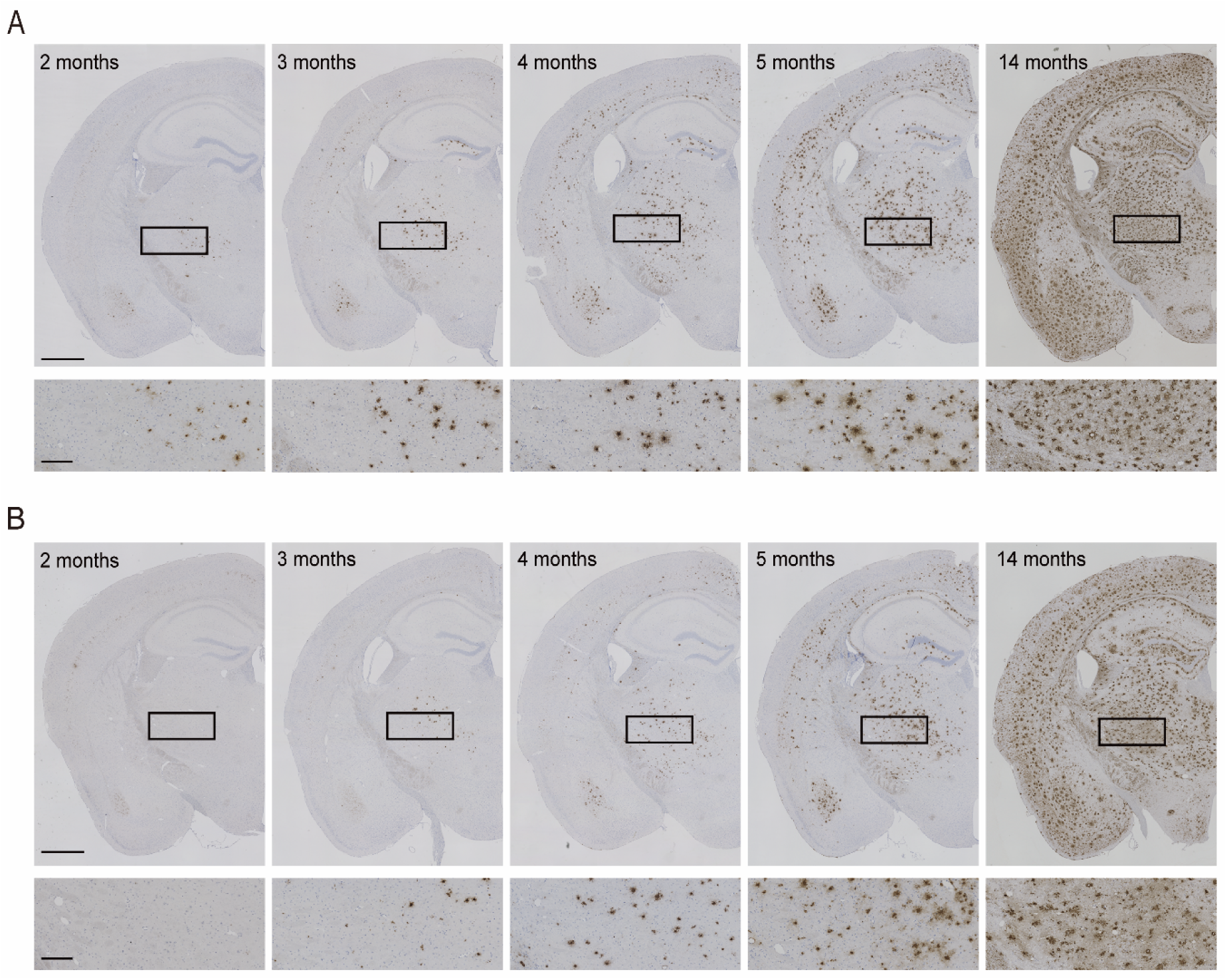
Monthly progression of Aβ deposition in preclinical 5XFAD mice. **(A and B)** Representative images of Aβ immunostaining using 4G8 antibody in coronal brain sections of female (A) and male (B) 5XFAD mice in the preclinical (2–5 months old) and advanced clinical (14 months old) stages. The lower panels are magnified images of the insets in the upper panels. Scale bars, 1 mm (upper panels), 250 µm (lower panels).

